# Demonstration of periodic and aperiodic EEG reliability between laboratory and clinic settings

**DOI:** 10.64898/2026.02.03.703614

**Authors:** Erin S.M. Matsuba, Haerin Chung, Alex Job Said, Margaret Norberg, Charles A. Nelson, Carol L. Wilkinson

**Affiliations:** Boston Children’s Hospital, 2 Brookline Place, Brookline, MA 02445, USA; Harvard Medical School, 25 Shattuck Street, Boston, MA 02115, USA

**Keywords:** EEG, reliability

## Abstract

**Objective:** To facilitate the scalability of EEG research, this paper compares the data quality and evaluates the absolute agreement of EEG features between laboratory and clinic settings.

**Methods:** Resting state EEG recordings were obtained from 36 participants (11 infants, 10 children, and 15 adults) from the waiting room of a primary care clinic and a laboratory. Intraclass correlation coefficients (ICC(2,1)) quantified the absolute agreement between laboratory and clinic settings for periodic power bands, alpha peak characteristics, and aperiodic components. The mean absolute difference (MAD) between laboratory and clinic recorded EEGs were calculated to describe signal consistency across settings.

**Results:** More components were rejected from clinic-recorded EEGs, though data quality otherwise did not differ between settings. The ICC (2,1) for all EEG measures were generally in the good-to-excellent range across ages and regions of interest. The MAD decreased with age and was largest in the alpha frequency range.

**Conclusions:** High quality EEG data can be collected from outpatient clinic settings among infants, children, and adults. There is high reliability in the parameterized periodic and aperiodic EEG features between laboratory and clinic settings.

**Significance:** Future research may collect EEG datasets from naturalistic settings with confidence in their reliability relative to laboratory recordings.

## Introduction

Electroencephalography (EEG) has been instrumental in our understanding of cognition and brain function (Cohen, 2014). More recently, EEG has emerged as a powerful tool in the effort to identify biomarkers for neurodevelopmental conditions (McPartland, 2016; Slater et al., 2022; Webb et al., 2023). EEG has many advantages that are well-suited for clinical utility; it is relatively inexpensive, non-invasive, and well-tolerated (Goodspeed et al., 2023; Webb et al., 2023). Resting-state EEG can be acquired in under five minutes and does not require task engagement, making it inclusive to most populations. Further, EEG equipment already exists in many clinical settings to support the diagnosis of medical conditions (e.g., epilepsy, sleep disorders) and EEG data collection can be integrated within existing patient visit structures (e.g., well-child visits; Conroy et al., 2025). Thus, the primary care setting is a logical and feasible location for the clinical implementation of EEG-based biomarkers, as well as a method to increase general access to EEG research participation. Even so, only few studies have assessed the potential of translating laboratory-based EEG findings into patient care settings. Ensuring the effective translation of EEG discoveries from basic science to clinical practice is essential for advancing healthcare quality, equity, and accessibility.

Significant efforts have been made to identify candidate EEG-based biomarkers for neurodevelopmental disorders. For instance, power across different frequency bands can predict autism outcomes (Bosl et al., 2018; Gabard-Durnam et al., 2019; Neo et al., 2023) and theta/beta ratios are associated with ADHD (Arns et al., 2013; Begum-Ali et al., 2022). Recent methodological advances in EEG have enabled a parametrization of absolute power to isolate the putative periodic and aperiodic spectra (Donoghue et al., 2020). This is crucial because conflating oscillatory (periodic) activity with non-oscillatory (aperiodic) activity may result in misattributions of important neurophysiological, cognitive, and behavioral results (Donoghue et al., 2020; Euler et al., 2024; Kałamała et al., 2024). Recent research suggests that components of aperiodic activity (i.e., 1/f curve, slope, and offset) are far from meaningless and may represent potential biomarkers for neurodevelopmental disorders (Arnett et al., 2024; Wilkinson et al., 2024). Thus, research that parameterizes these distinct components is important and timely such that we may truly understand the underlying neurophysiology.

While significant research has focused on identifying EEG biomarkers, these studies have assessed whether EEG features correlate with symptoms (i.e., construct validity) or predict future diagnoses (i.e., predictive validity). However, clinical utility depends on both validity and reliability, where validity refers to the correctness against the ground truth signal accuracy, and reliability refers to the consistency across multiple measurements and settings (Cook & Beckman, 2006; Vetter & Cubbin, 2019). Positively, research on test-retest reliability of resting state EEG features in laboratory settings has found generally acceptable results among children and adults across periodic power bands, as well as aperiodic slope and offset (Levin et al., 2020; Lopez et al., 2023; Webb et al., 2023).

Less research has evaluated whether resting state EEG measures are generalizable beyond a highly controlled laboratory environment. Recent technological (e.g., mobile, portable, and wireless EEG systems) and methodological advances (e.g., standardized pre-processing pipelines, independent component analysis; ICA) have coincided to enable cognitive neuroscience research in real-world settings (Ladouce et al., 2017). However, few studies focus on the ecological validity of EEG beyond the laboratory and fewer still have directly compared EEG signals across settings. Mikkelsen et al. (2021) found that EEG data quality was similar between laboratory and home settings among adults (11% of epochs rejected in laboratory; 7% of epochs rejected at home), with some evidence to suggest that inter- and intra-subject variability was greater than that of location. Recently, Dickinson et al. (2025) evaluated the reliability of absolute EEG power bands across laboratory and community settings among a small sample (*n* = 10) of young children with developmental concerns. Despite the use of different EEG caps by setting, intraclass correlations found good absolute agreement between laboratory and home settings for beta and gamma absolute power across all regions of interest (i.e., frontal, central, parietal, and occipital), though delta, theta, and alpha power ranged from poor-to-moderate agreement by region.

While these studies provide preliminary data to support the reliability of EEG signals in naturalistic settings, research has yet to quantify the test-retest reliability of the parameterized periodic and aperiodic spectra. If EEG is to be used as a screening measure for the early identification of neurodevelopmental disorders (and increase general EEG research participation), it would be ideal to collect data in familiar and accessible settings. As such, the primary aim of this study was to evaluate the reliability of resting state EEG features between a laboratory and an outpatient primary care setting.

## Methods

### Participants

Thirty-six participants participated in this study. This included 11 infants (mean age = 5.7 months; SD = 2.5 months), 10 children (mean age = 5.5 years; SD = 4.0 years), and 15 adults (mean age = 28.1 years; SD = 5.8 years). Chronological age was calculated as the age at the date of the laboratory EEG visit. All participants had no known history of prenatal or postnatal medical or neurological problems, and no known genetic disorders. One infant fell asleep during the clinic EEG recording and was excluded from subsequent analyses. Two additional participants did not meet the EEG data quality requirements. Thus, the final sample for analysis consisted of 33 participants (*n* = 10 infants, *n* = 10 children, *n* = 13 adults).

### Experimental Paradigm

Each participant completed two 5-minute EEG recordings, one at the laboratory and one at the clinic, both using a 128-channel Hydrocel Geodesic Sensor Nets (Electrical Geodesics, Inc., Eugene, OR) connected to a NetAmps 400 amplifier, sampled at 1000 Hz with a 0.1 Hz high-pass analog filter, and online re-referencing to the vertex (Cz electrode) through NetStation software. Impedances were maintained below 50KΩ. Participants were guided to remain calm while they watched a non-task-related video of toys. Trained research assistants were present in both settings and provided a quiet toy (e.g., a rattle) to the infant or child if they became distressed. Recordings were paused during any periods of increased motor activity or emotional dysregulation (e.g., crying, fussing, coughing). Infants were always seated on their caregiver’s lap, while children and adults sat independently. On average, the laboratory and clinic visits were 3.62 days apart (range = 0-20 days; SD = 4.87).

The laboratory-based EEG was collected from a shielded, sound-attenuated room designed to block electrical fields, electromagnetic interference, and radio frequencies. The clinic-based EEG was recorded in an outpatient primary care clinic using a similar EEG system with the amplifier and computer secured on a mobile cart (Figure 1). Recordings were collected in a quiet area of the clinic adjacent to the waiting room. Light, sound, and electrical noise measures were collected from both settings over a series of 5 days using Gauss, decibel, and light meters. As expected, the laboratory setting had lower light, sound, and electrical noise levels, as well as less variability across measurements (Supplementary Materials Table 1).

**Table 1.**
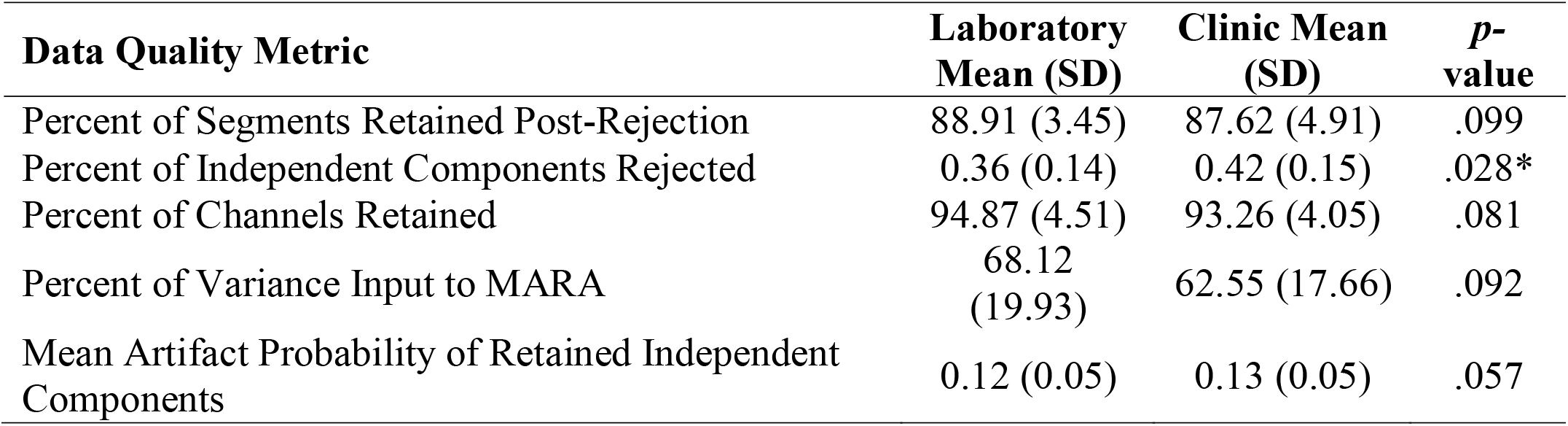
Comparison of data quality measures between laboratory and clinic settings.

**Figure 1.**
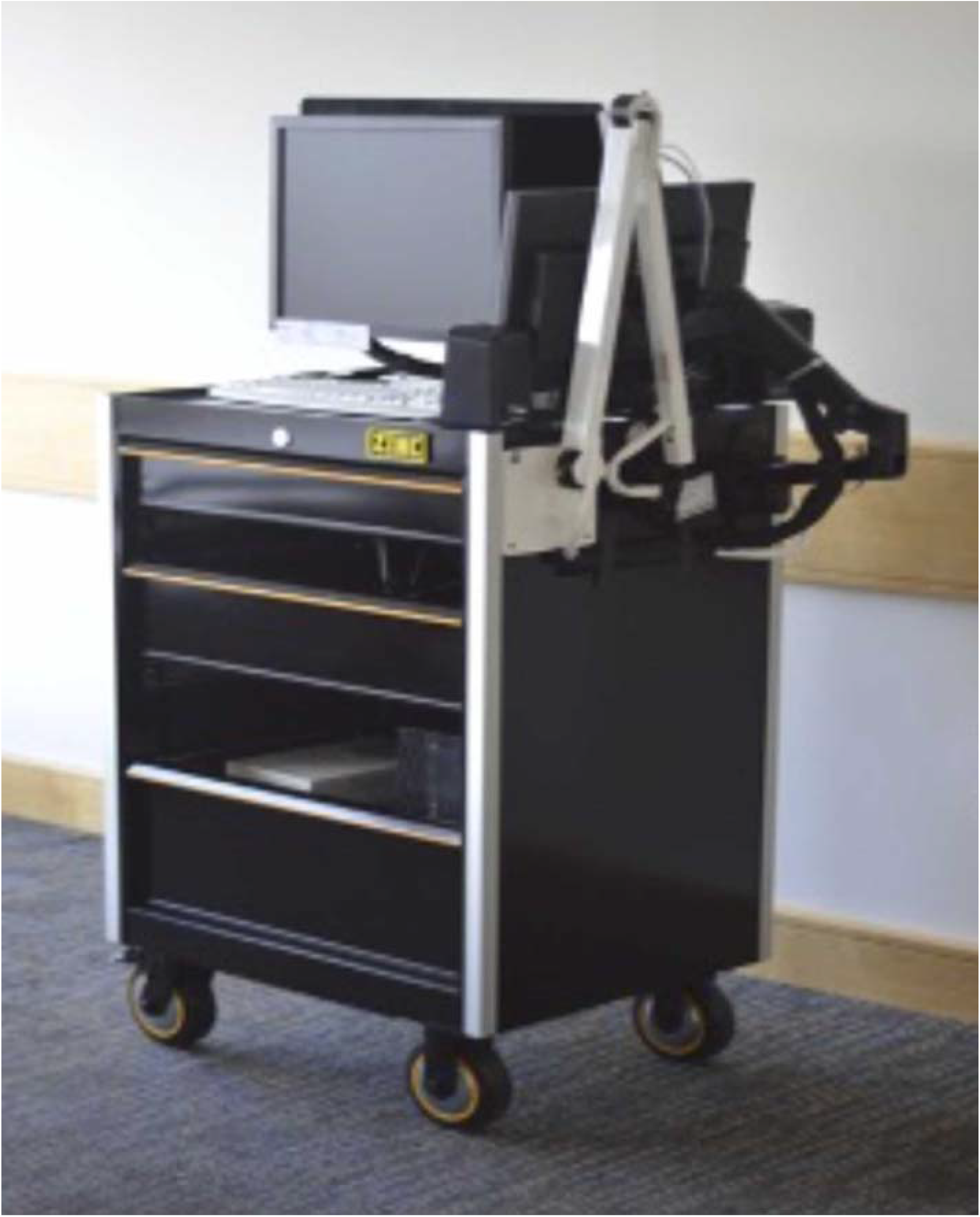
Mobile EEG Cart in the outpatient primary care clinic.

### EEG pre-processing

EEG data were pre-processed using the standardized Batch EEG Automated Processing Platform (BEAPP; Levin et al., 2018), in combination with Harvard Automated Processing Pipeline for Electroencephalography (HAPPE; Gabard-Durnam et al., 2018). In order to optimize artifact rejection performance, a spatially distributed subset of 36 channels providing whole-head coverage were processed through HAPPE (electrodes 4, 9, 11, 13, 19, 22, 24, 28, 33, 36, 37, 41, 45, 47, 52, 55, 58, 62, 65, 67, 70, 75, 77, 83, 87, 90, 92, 96, 98, 103, 104, 108, 112, 117, 122, and 124). EEG data was filtered with a 1 Hz high-pass filter, 100 Hz low-pass filter, and notch filter with CleanLine (Mullen, 2012) to remove 60 Hz line noise.

EEG data were then resampled with interpolation to 250□Hz (resampling was performed after filtering to avoid aliasing higher frequencies when resampling) in preparation for ICA. Artifacts were rejected (e.g., eye blinks, movement, and muscle activity) first through wavelet-enhanced ICA (wICA), followed by automated ICA rejection using MARA (Winkler et al., 2011). Post-artifact rejection, any channels removed during the bad channel rejection were interpolated through spherical interpolation to reduce spatial bias in re-referencing. The EEG data were then re-referenced to the average reference, detrended to the signal mean, and segmented into 2 second windows. Additional segments were then rejected using HAPPE’s amplitude and joint probability criteria.

EEG recordings were rejected if the pre-processed and cleaned EEG data had fewer than 20 segments (40 seconds of total EEG), the number of good channels were below 80%, the mean or median retained artifact probability was above 30%, the percent independent components rejected as artifact was above 80%, or the percent variance retained after artifact removal was under 25%. Following this criteria, one adult EEG from the clinic was removed because more than 80% of their components rejected by wICA and one adult EEG from the laboratory had too little variability input to MARA.

### EEG spectral decomposition and parameterization analysis

The power spectral density (PSD) was calculated using multitaper spectral analysis with three orthogonal tapers. For each electrode, the PSD was averaged across segments, and then additionally averaged over electrodes from regions of interest (frontal: 4, 11, 19, 24, 28, 117, 124; central: 13, 36, 37, 55, 87, 104, 112; posterior: 62, 67, 77, 70, 75, 83; temporal: 45, 41, 47, 52, 108, 103, 98, 92; Supplemental Figure 1). The PSD was then further analyzed using a modified version of SpecParam v1.0.0 (also known as FOOOF, https://github.com/fooof-tools/fooof; in Python v3.6.8) to model periodic and aperiodic components of the power spectra. SpecParam was set to the fixed mode (without a spectral knee), with peak_width_limits set to [0.5, 18.0], max_n_peaks = 7, and peak_threshold = 2. To characterize the aperiodic exponent (i.e., the slope of the spectra), we used χ in the 1/fχ model fit. Aperiodic offset was defined as the aperiodic power at 2.5 Hz. Further analyses were subsequently restricted to 2.5-50 Hz given elevated error between 2-2.5 and 50-55 Hz. Participants with a poor SpecParam fit (defined as R^2^< 0.97) were excluded from further analyses to more confidently compare the parameterized spectra. This resulted in the removal of 8 participants across ROIs (central *n* = 1 laboratory and 1 clinic EEG, frontal *n* = 1 clinic EEG, posterior *n* = 1 laboratory EEG, temporal *n* = 2 laboratory EEGs, whole *n* = 3 laboratory EEGs). Final SpecParam fit ranged from 0.996-0.999 and was comparable across settings, ROIs, and age groups (See Supplementary Table 2). Power for each band was calculated by taking the integral of the periodic spectra between the following frequency ranges: theta (4-6 Hz), low alpha (6-9 Hz), high alpha (9-12 Hz), low beta (12-20 Hz), high beta (20-30 Hz), gamma (30-45 Hz), and total power (2.5-50 Hz). Alpha peak amplitude was calculated as the maximum power from 4-12 Hz and alpha peak frequency was the corresponding frequency (Hz).

### Statistical Analysis

EEG data quality measures across settings were submitted to a paired samples t-test. Intraclass correlations using a two-way random effects model (ICC(2,1)) were used to estimate absolute agreement between EEG data from laboratory and clinic settings for the canonical periodic frequency bands of low alpha, high alpha, low beta, high beta, delta, gamma, and theta, peak characteristics of alpha frequency and amplitude, as well as aperiodic offset and slope (Figure 2).

**Figure 2.**
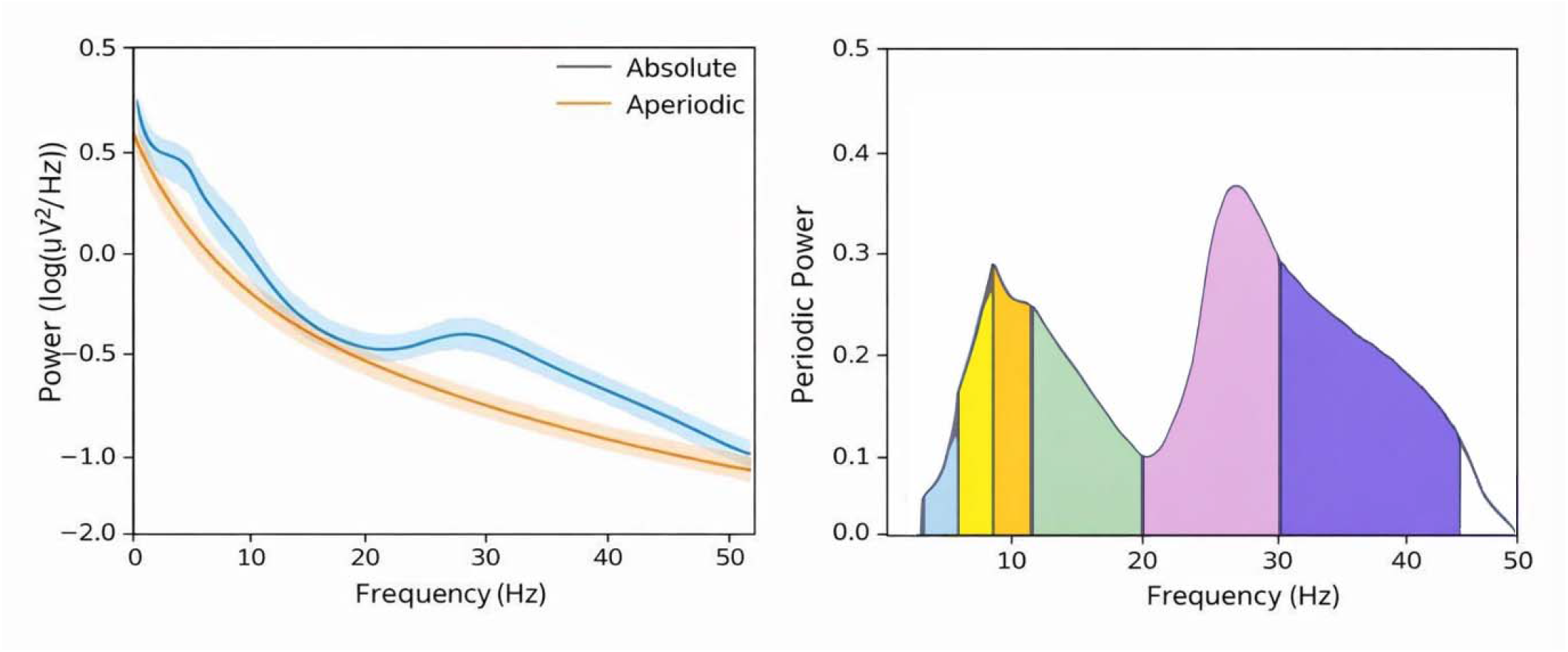
Example absolute and aperiodic spectra with aperiodic offset and slope (left) and periodic spectra with canonical frequency bands of theta (4-6 Hz), low alpha (6-9 Hz), high alpha (9-12 Hz), low beta (12-20 Hz), high beta (20-30 Hz), and gamma (30-45 Hz) (right).

ICC values were interpreted using typical descriptors in which values less than 0.4 are considered poor, 0.4-0.59 are fair, 0.6-0.74 are good; 0.75-1 are considered excellent (Cicchetti, 1994). To further visualize and describe differences between laboratory and clinic settings by age group, the mean absolute difference (MAD) between each participant’s laboratory and clinic absolute, periodic, and aperiodic spectra were calculated (laboratory spectra– clinic spectra power). Because spectra were log-transformed, the MAD represents the mean absolute log-difference, which corresponds to a relative difference (log-ratio) between laboratory and clinic recordings. To our knowledge, there are no established guidelines on the acceptable MAD range for absolute, periodic, or aperiodic spectra, and these measures are not directly comparable to each other due to their differing units. Accordingly, the MAD were used qualitatively to describe patterns across ROIs and age groups.

## Results

### Data Quality across Settings

Paired sample t-tests revealed that data quality did not differ between laboratory and clinic settings (see Table 1), with the exception of significantly more independent components (i.e., unique source signals) rejected in the clinic compared to laboratory setting (*t*(33) = -2.3, *p* = 0.028).

### Intraclass Correlations between Laboratory and Clinic EEG Spectra

Parameterized power from each of the canonical frequency bands was consistent with previous research (see Supplementary Materials Table 3). Across participants, the power spectra appeared similar across laboratory and clinic settings (Figure 3). Two-way random effects intraclass correlations (ICC(2,1)) for absolute agreement were computed to assess the reliability of resting-state EEG features across common ROIs. Overall, ICCs revealed that the absolute agreement between laboratory and clinic settings was in the good-to-excellent range for all periodic and aperiodic EEG spectral features (Figure 4). Reliability was highest across ROIs for high alpha (range = 0.79-0.82), low beta (range = 0.86-0.88), high beta (0.83-0.87), gamma (range = 0.76-0.87), broad alpha peak frequency (range = 0.75-0.94), and aperiodic offset (range = 0.75-0.88). Alpha peak amplitude demonstrated the lowest absolute agreement (range = 0.59-0.73), though was still in the fair-to-good range. Across regions, central and frontal ROIs generally exhibited the highest reliability, whereas temporal and posterior ROIs were slightly lower. A full summary of the ICC results may be found in Supplementary Materials Table 4.

**Figure 3.**
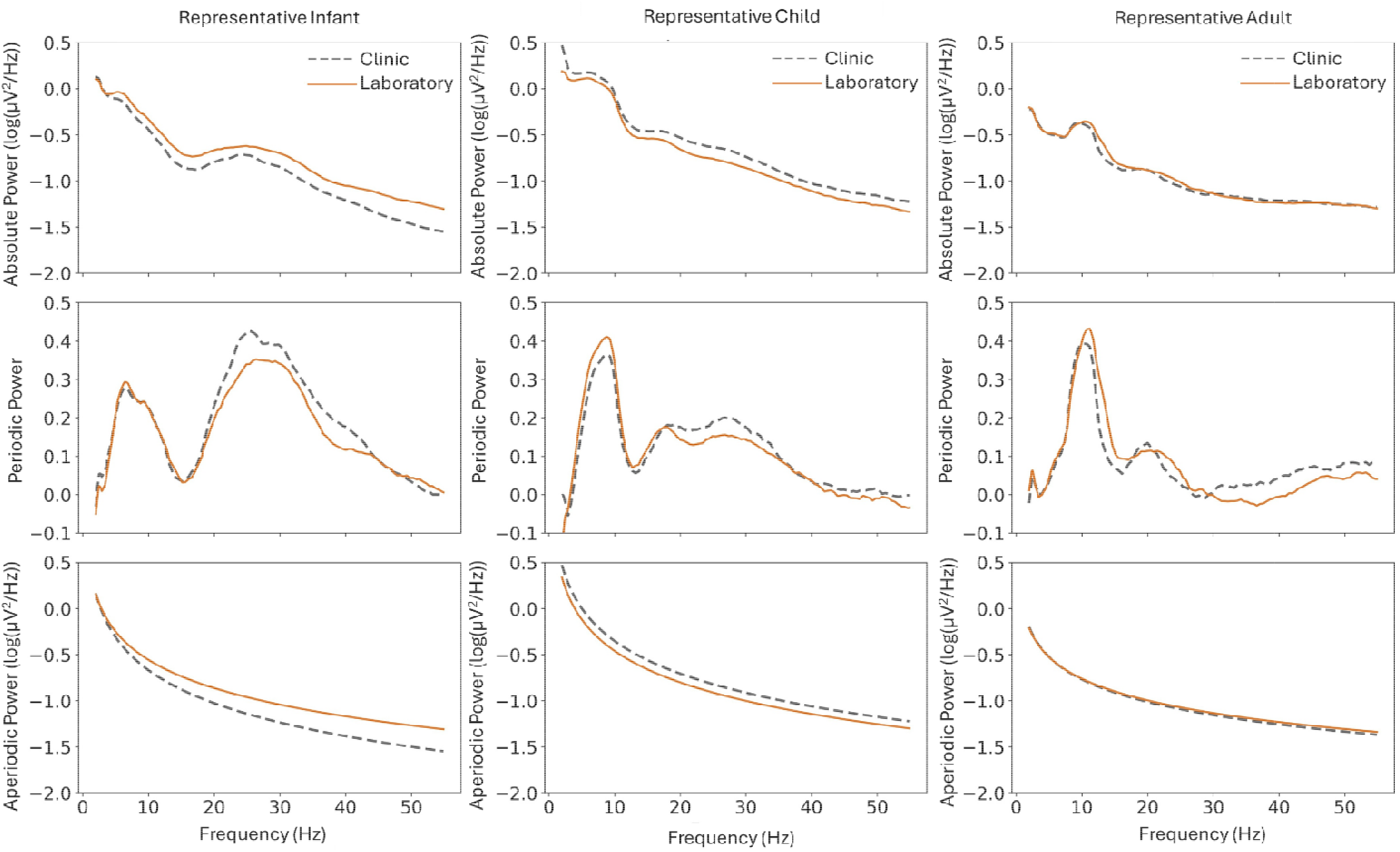
Representative participants’ absolute (top), periodic (middle), and aperiodic (bottom) spectral power across clinic and laboratory settings.

**Figure 4.**
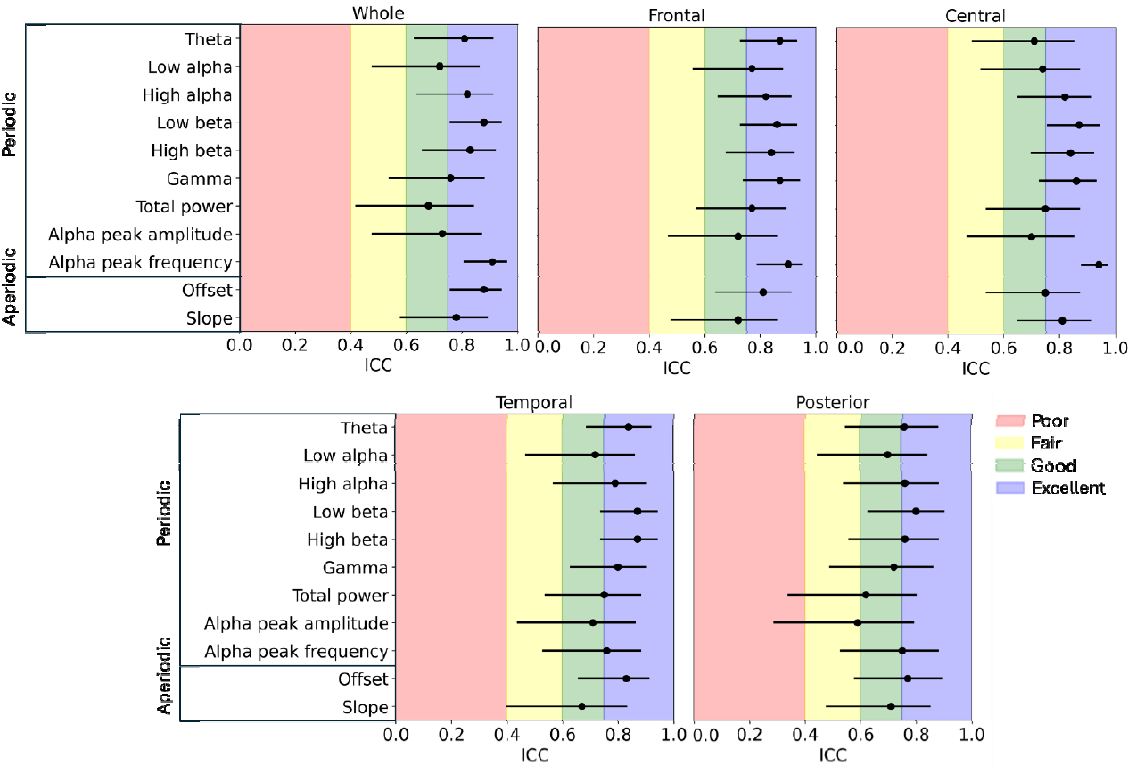
Intraclass correlations (ICC(2,1)) between laboratory and clinic settings for periodic and aperiodic components across regions of interest (ROI).

### Visualizing Differences between Laboratory and Clinic EEG Spectra

The MADs by spectral component, ROI, and age group are presented to further describe and visualize the EEG spectra differences across laboratory and clinic settings (Supplementary Table 5). Overall, the MAD showed age-related improvements across most spectral features, with infants and children exhibiting more discrepancies compared to adults. Regarding ROIs, the frontal and central electrodes consistently yielded the lowest MADs, while posterior and temporal regions showed slightly more variability, particularly among infants and children. While no previous interpretive guidelines exist for the MAD of two EEGs in distinct locations, Ostlund et al. (2022) indicates that a mean absolute error (MAE) between 0.025-0.1 indicates acceptable spectral fit between original and SpecParam-modeled spectra. The absolute spectral power of laboratory relative to clinic recorded EEGs were similar to this range for adults (range 0.10-0.15), though higher for children (range = 0.10 – 0.17) and infants (range = 0.12-0.19). The relative difference between laboratory and clinic EEG for periodic power was similar across ROIs and age groups (range = 0.04-0.08). Visual analysis indicates that across most ROIs, the periodic spectra MAD was largest around 10 Hz (Figure 5). The MAD for aperiodic features was generally lower among adults (range = 0.09-0.15). For infants and children, the aperiodic spectra MAD was more variable (range = 0.11-0.22) and tended to increase with frequency across ROIs (Figure 5).

**Figure 5.**
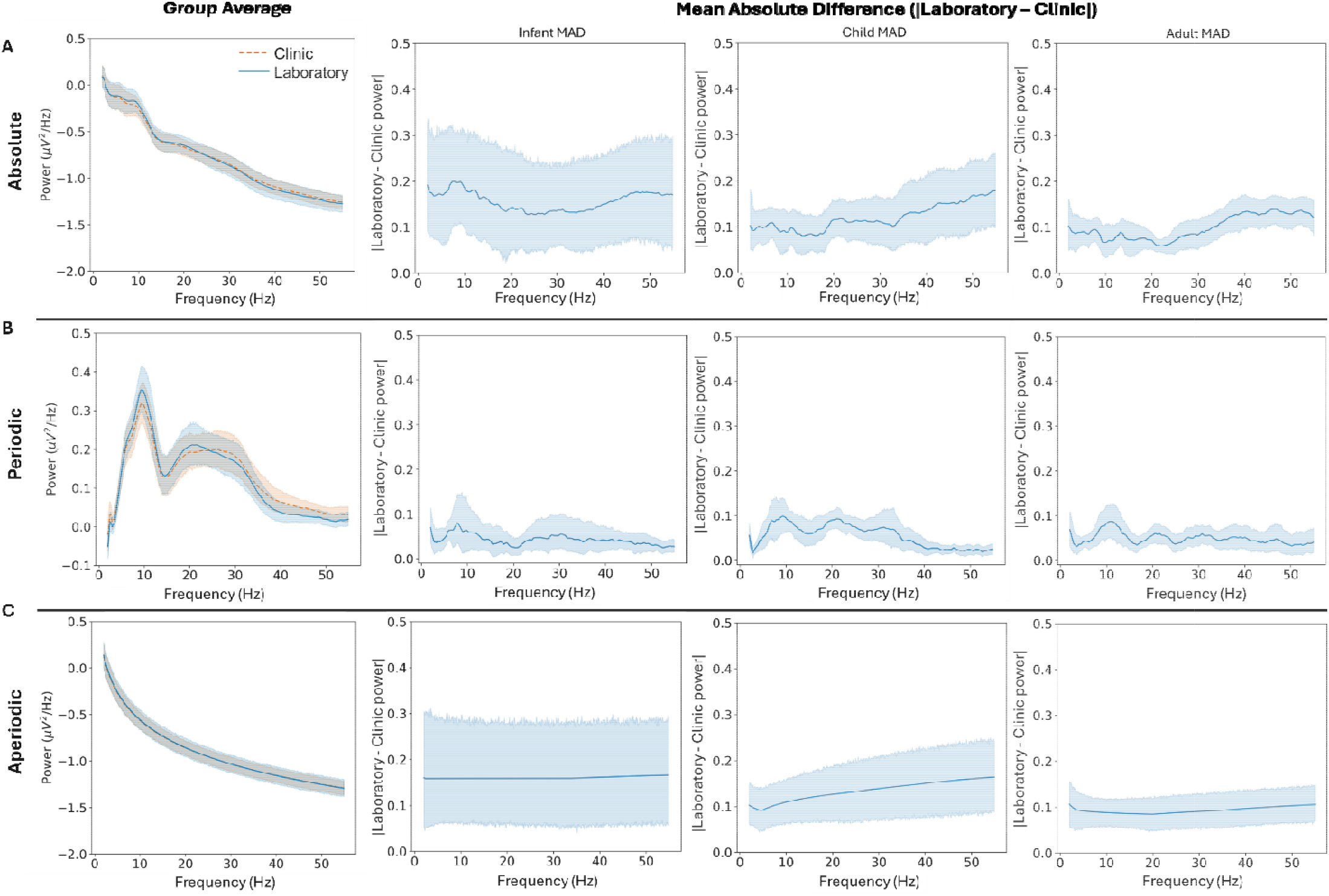
Mean absolute difference (MAD) between the laboratory and clinic EEG across absolute (top), periodic (middle), and aperiodic (bottom) spectra among infants (left), children (middle), and adults (right) from the whole region of interest.

## Discussion

The present study sought to evaluate the reliability of collecting parameterized resting-state EEG features in a controlled laboratory environment and a real-life outpatient clinic setting among infants, children, and adults. In so doing, this study addresses a gap in the literature regarding the impact of location on the reliability of the EEG signal, which is crucial for the implementation of EEG-based candidate biomarkers of neurodevelopmental disorders.

High quality data from EEGs recorded in the clinic is a necessary prerequisite to determine whether the signals are reliable across settings. As described above, lessons from previous research have resulted in effective strategies to reduce noise during data collection (e.g., pausing the recording if participants moved or cried, maintaining impedances below 50KΩ) and pre-processing for artifact removal (e.g., wICA and MARA). We found that final data quality metrics between laboratory and clinic settings did not differ, despite some evidence to suggest more noise in the initial clinic-based recording. That is, while there were no differences between laboratory and clinic recorded EEGs regarding the percentage of rejected segments or bad channels, significantly more components were rejected from data recorded in the clinic.However, the mean probability of artifact in retained components did not differ. Together, this indicates that while real-world clinic settings may introduce more artifact-based components (as expected in locations with more light, sound, and electrical noise), standardized pre-processing pipelines are well-equipped to identify and remove those artifacts to produce high quality EEG data for subsequent analyses (Levin et al., 2018; Gabard-Durnam et al., 2018). Thus, despite a greater burden of artifact rejection in real-world settings, EEG data quality collected from the primary care clinic may be comparable to that of highly controlled laboratory settings.

The test-retest reliability between laboratory and clinic settings for most periodic EEG features (i.e., periodic high alpha, low beta, high beta, and gamma power, as well as alpha peak frequency) were in the excellent range across all ages and ROIs. Theta power, low alpha power, and alpha peak amplitude demonstrated generally good reliability (though see alpha peak amplitude in the posterior ROI). Overall, this indicates that there is a high degree of reliability between EEGs collected in controlled laboratory and primary care clinic settings. Agreement was higher in this study compared to previous assessments of EEGs between laboratory and community settings (Dickinson et al., 2025). Certainly, several key methodological differences may explain these differences (e.g., sample size and age range, different pre-processing procedures). Perhaps most saliently, agreement in previous work was complicated by use of different EEG systems by setting (Dickinson et al., 2025), suggesting that consistency in EEG cap and recording system is an important factor for test-retest reliability across locations. Further, here we present novel findings on the reliability of parameterized components, namely periodic and aperiodic spectra, recorded from laboratory and clinic settings. Previous studies report poor reliability in low frequency absolute power bands (i.e., delta and theta; Dickinson et al., 2025), which may be driven by decreased agreement in the aperiodic spectra that disproportionately impacts lower frequencies. In the present study, we see increased relative differences in aperiodic power across settings, particularly among the age range of Dickinson et al. (2025). Thus, the implementation of parameterization algorithms (i.e., SpecParam; Donoghue et al., 2020) may enable more precise measurement to isolate and understand how distinct components of the absolute power spectra change across different recordings and locations.

To further analyze the periodic spectra, the MAD between periodic laboratory and clinic spectra was computed and observed to be the largest in the alpha range (6-12 Hz). Alpha power is sensitive to light, with increased alpha power observed in dark settings. Given that lux levels were higher in the clinic, it is perhaps unsurprising that reductions in alpha power were observed in the clinic-based EEGs (Berger et al., 1933). Further, as alpha power is also an indicator for arousal or engagement, this may suggest greater activation, sensory processing, and attention to external stimuli in the clinic. Though intraclass correlations were in the good-to-excellent range for alpha power and alpha peak frequency, this may suggest a need to evaluate any alpha-based candidate biomarker in different lightings to maximize precision and generalizability to real world settings. Notably, differences in aperiodic fit between settings may impact the subsequent parameterized periodic spectra (Donoghue et al., 2020). However, a *posthoc* correlation of the periodic alpha MAD and aperiodic slope MAD was not significant (*r* = -0.07, *p* = 0.734), suggesting that differences cannot be explained by aperiodic fit alone. Further, differences in alpha between laboratory and clinic settings were appreciable in the periodic MAD but not the absolute or aperiodic MAD, further demonstrating the importance of methodological innovations that enable a deeper understanding of the underlying neurophysiology.

Regarding aperiodic features, reliability between laboratory and clinic settings was excellent for offset and good for slope. Previous research has similarly observed higher reliability for offset compared to slope (Levin et al., 2020). Here, offset was re-calculated at 2.5 Hz to account for poor SpecParam model fit from 0-2.5 Hz, which may also have benefited reliability across recordings. In contrast, slope was derived from the overall morphology of the power spectrum and may have been variably modulated by many factors (e.g., state of the participant, movement). Indeed, visual inspection of the MAD suggests that while artifacts were similarly identified and removed by pre-processing across settings, remainder traces of muscle artifact may have disproportionately impacted higher frequencies (Figure 5) and decreased slope reliability. Nonetheless, by demonstrating good-to-excellent reliability between laboratory and clinic settings for aperiodic components, we hope to contribute to the growing literature that aperiodic activity represents stable and meaningful data that warrants further research.

### Limitations and Future Directions

A fundamental aspect of our study was to assess whether EEG signals collected in a real-world primary care clinic with variability in environmental conditions (e.g., electrical, auditory, and light noise) were similar to that collected in the laboratory. By design, some recordings at the clinic more closely resembled a controlled and quiet laboratory, while others had ample background or electrical noise that may have unevenly influenced the EEG signal and participant behavior. Additionally, despite our sample size being larger than prior studies, we were not able to statistically assess differences in test-retest reliability between age groups. Lastly, there are no standardized guidelines for evaluating the MAD across settings, which limited our ability to compare and interpret our results. Thus, future studies with large, diverse, and repeated samples may further increase confidence that resting state EEG features are reliable across laboratory and clinic settings.

## Conclusions

In conclusion, the present study sought to improve the scalability and inclusivity of EEG in early developmental research by comparing EEGs obtained in the clinic and laboratory settings. To that end, we leveraged standardized and openly-available pre-processing pipelines (i.e., BEAPP, HAPPE) combined with spectral decomposition into aperiodic and periodic components (i.e., SpecParam) to demonstrate that periodic power bands and aperiodic components are robust measures that can be collected reliably from a primary care clinic.Additionally, separately quantifying the periodic and aperiodic spectra allowed for greater precision and understanding of the agreement between neurophysiological signatures by location, which highlights the value and need for greater research and attention to these methodological considerations. Our findings indicate the capacity to collect high quality data from real-world clinics (even among infants and children) and suggest that researchers may combine datasets collected from different locations. Lastly, this novel demonstration of ecological validity supports the implementation readiness of EEG-based biomarkers to outpatient settings, which could ultimately yield earlier identification of neurodevelopmental conditions in convenient, accessible clinic settings.

## Notes

**Conflict of Interest Statement:** None of the authors have potential conflicts of interest to be disclosed.

**Funding:** This work was generously supported by the Eagles Autism Foundation.

### Competing Interest Statement

The authors have declared no competing interest.

