## Supplemental Materials for "Demonstration of periodic and aperiodic EEG reliability between laboratory and clinic settings"

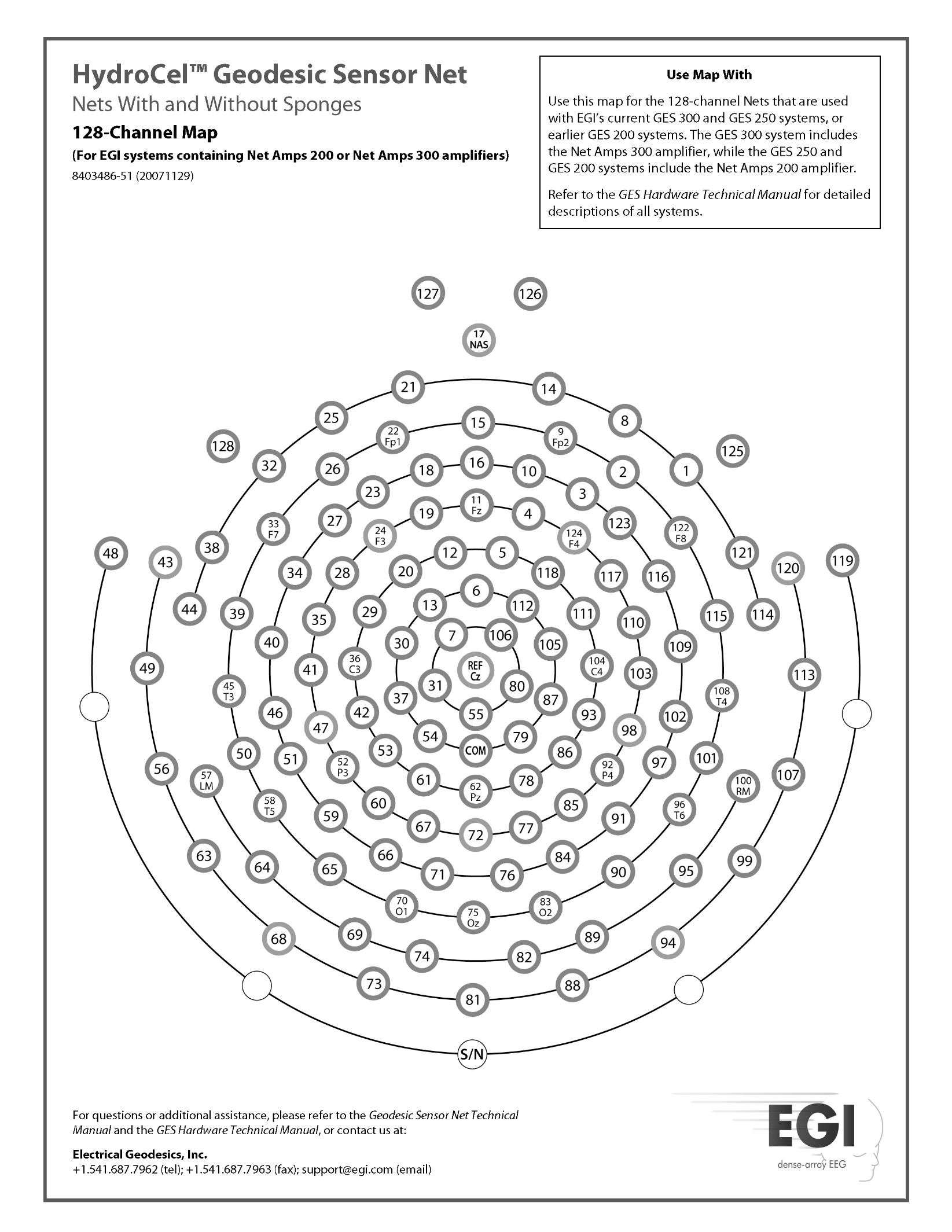
Supplementary Materials

Fig. S2. Electrode layout: 128-channel Hydrocel Geodesic Sensor Net. Red, green, skyblue, and yellow circles denote electrodes included in ICA and MARA steps of pre-processing. Electrodes averaged for frontal (red), central (green), and posterior (yellow) regions of interest.

| **Table 1. Summary of light, sound, and electrical noise across laboratory and clinical settings** | | |
| --- | --- | --- |
|  | Clinic (SD) | Laboratory (SD) |
| Light Level (lux) | 170.64 (22.97) | 30 (2) |
| Electrical Noise (Gauss) | 0.83 (0.75) | 0.34 (0.24) |
| Sound Level (dB) | 52.54 (4.62) | 33 (3.01) |

| **Table 2.** SpecParam model fit by region of interest, age group, and location (mean (SD)) | | | | | | |
| --- | --- | --- | --- | --- | --- | --- |
| **Age Group** | **Location** | **Central** | **Frontal** | **Posterior** | **Temporal** | **Whole** |
| Infants | Clinic | 0.998 (0.003) | 0.998 (0.001) | 0.999 (0.001) | 0.998 (0.002) | 0.999 (0.001) |
|  | Laboratory | 0.999 (0.002) | 0.998 (0.002) | 0.997 (0.006) | 0.998 (0.001) | 0.999 (0.001) |
| Children | Clinic | 0.998 (0.002) | 0.998 (0.001) | 0.995 (0.006) | 0.998 (0.001) | 0.998 (0.002) |
|  | Laboratory | 0.996 (0.008) | 0.997 (0.005) | 0.996 (0.007) | 0.997 (0.003) | 0.998 (0.002) |
| Adults | Clinic | 0.995 (0.005) | 0.997 (0.002) | 0.995 (0.004) | 0.993 (0.008) | 0.994 (0.005) |
|  | Laboratory | 0.993 (0.008) | 0.997 (0.003) | 0.996 (0.003) | 0.994 (0.007) | 0.995 (0.004) |

| **Table 3.** Descriptive statistics for periodic and aperiodic spectra features (mean (SD)) | | | | | | |
| --- | --- | --- | --- | --- | --- | --- |
|  |  | Whole | Frontal | Central | Temporal | Posterior |
| Theta | Laboratory | 0.26 (0.17) | 0.30 (0.19) | 0.31 (0.18) | 0.31 (0.20) | 0.24 (0.18) |
|  | Clinic | 0.26 (0.18) | 0.31 (0.18) | 0.34 (0.17) | 0.31 (0.20) | 0.23 (0.19) |
| Low alpha | Laboratory | 0.78 (0.33) | 0.86 (0.34) | 0.96 (0.42) | 0.89 (0.37) | 0.82 (0.39) |
|  | Clinic | 0.72 (0.36) | 0.85 (0.33) | 0.96 (0.41) | 0.78 (0.38) | 0.72 (0.40) |
| High alpha | Laboratory | 0.93 (0.52) | 1.00 (0.52) | 1.18 (0.54) | 1.05 (0.60) | 1.05 (0.49) |
|  | Clinic | 0.83 (0.42) | 0.95 (0.44) | 1.12 (0.49) | 0.90 (0.51) | 0.94 (0.40) |
| Low beta | Laboratory | 1.36 (1.08) | 1.64 (1.11) | 1.83 (1.07) | 1.66 (1.18) | 1.47 (1.00) |
|  | Clinic | 1.27 (1.03) | 1.63 (1.01) | 1.86 (1.06) | 1.47 (1.14) | 1.37 (0.98) |
| High beta | Laboratory | 1.92 (1.13) | 2.70 (1.36) | 2.66 (1.37) | 2.42 (1.30) | 1.61 (1.15) |
|  | Clinic | 1.93 (1.11) | 2.74 (1.35) | 2.83 (1.46) | 2.31 (1.26) | 1.53 (1.22) |
| Gamma | Laboratory | 1.11 (1.17) | 1.51 (1.40) | 1.75 (1.57) | 1.63 (1.39) | 1.01 (1.05) |
|  | Clinic | 1.40 (1.25) | 1.70 (1.43) | 1.99 (1.69) | 1.77 (1.39) | 1.13 (1.23) |
| Total power | Laboratory | 6.62 (2.81) | 8.37 (3.40) | 9.05 (3.36) | 8.32 (3.30) | 6.48 (2.89) |
|  | Clinic | 6.78 (2.65) | 8.56 (2.98) | 9.54 (3.61) | 7.96 (3.06) | 6.28 (3.30) |
| Offset | Laboratory | 0.04 (0.35) | 0.02 (0.28) | -0.15 (0.42) | -0.12 (0.36) | 0.10 (0.39) |
|  | Clinic | 0.03 (0.34) | -0.02 (0.26) | -0.21 (0.33) | -0.12 (0.34) | 0.16 (0.32) |
| Slope | Laboratory | 1.00 (0.16) | 1.07 (0.16) | 1.12 (0.19) | 1.00 (0.17) | 1.04 (0.18) |
|  | Clinic | 0.98 (0.21) | 1.08 (0.18) | 1.10 (0.19) | 0.97 (0.19) | 1.07 (0.23) |

| **Table 4.** Intraclass Correlations (2,1) for periodic and aperiodic spectra features ICC (2,1) [95% CI] | | | | | |
| --- | --- | --- | --- | --- | --- |
|  | Whole | Frontal | Central | Temporal | Posterior |
| Canonical frequency bands of periodic spectra | | | | | |
| Theta | 0.81 [0.63, 0.91] | 0.87 [0.73, 0.93] | 0.71 [0.49, 0.85] | 0.84 [0.69, 0.92] | 0.76 [0.55, 0.88] |
| Low alpha | 0.72 [0.48, 0.86] | 0.77 [0.56, 0.88] | 0.74 [0.52, 0.87] | 0.72 [0.47, 0.86] | 0.70 [0.45, 0.84] |
| High alpha | 0.82 [0.64, 0.91] | 0.82 [0.65, 0.91] | 0.82 [0.65, 0.91] | 0.79 [0.57, 0.90] | 0.76 [0.54, 0.88] |
| Low beta | 0.88 [0.76, 0.94] | 0.86 [0.73, 0.93] | 0.87 [0.76, 0.94] | 0.87 [0.74, 0.94] | 0.80 [0.63, 0.90] |
| High beta | 0.83 [0.66, 0.92] | 0.84 [0.68, 0.92] | 0.84 [0.70, 0.92] | 0.87 [0.74, 0.94] | 0.76 [0.56, 0.88] |
| Gamma | 0.76 [0.54, 0.88] | 0.87 [0.74, 0.94] | 0.86 [0.73, 0.93] | 0.80 [0.63, 0.90] | 0.72 [0.49, 0.86] |
| Total power | 0.68 [0.42, 0.84] | 0.77 [0.57, 0.89] | 0.75 [0.54, 0.87] | 0.75 [0.54, 0.88] | 0.62 [0.34, 0.80] |
| Periodic Spectra Peak Characteristics | | | | | |
| Alpha Peak Amplitude | 0.73 [0.48, 0.87] | 0.72 [0.47, 0.86] | 0.70 [0.47, 0.85] | 0.71 [0.44, 0.86] | 0.59 [0.29, 0.79] |
| Alpha Peak Frequency | 0.91 [0.81, 0.96] | 0.90 [0.79, 0.95] | 0.94 [0.88, 0.97] | 0.76 [0.53, 0.88] | 0.75 [0.53, 0.88] |
| Aperiodic Spectral Features | | | | | |
| Offset | 0.88 [0.76, 0.94] | 0.81 [0.64, 0.91] | 0.75 [0.54, 0.87] | 0.83 [0.66, 0.91] | 0.77 [0.58, 0.89] |
| Slope | 0.78 [0.58, 0.89] | 0.72 [0.48, 0.86] | 0.81 [0.65, 0.91] | 0.67 [0.40, 0.83] | 0.71 [0.48, 0.85] |

| **Table 5.** MAD between clinic and lab by region of interest by age group (mean (SD)) | | | | | |
| --- | --- | --- | --- | --- | --- |
| Absolute Spectrum (mv^2^/Hz) | **Central** | **Frontal** | **Posterior** | **Temporal** | **Whole** |
| Infants | 0.19 (0.29) | 0.12 (0.12) | 0.16 (0.13) | 0.14 (0.12) | 0.16 (0.16) |
| Children | 0.10 (0.07) | 0.11 (0.06) | 0.17 (0.14) | 0.17 (0.09) | 0.12 (0.08) |
| Adults | 0.10 (0.05) | 0.10 (0.04) | 0.14 (0.13) | 0.15 (0.06) | 0.10 (0.02) |
| Periodic Average (mv^2^/Hz) |  |  |  |  |  |
| Infants | 0.08 (0.05) | 0.06 (0.02) | 0.05 (0.03) | 0.05 (0.02) | 0.04 (0.03) |
| Children | 0.04 (0.02) | 0.05 (0.02) | 0.07 (0.04) | 0.05 (0.02) | 0.05 (0.03) |
| Adults | 0.05 (0.03) | 0.05 (0.02) | 0.06 (0.04) | 0.07 (0.04) | 0.05 (0.02) |
| Aperiodic (mv^2^/Hz) |  |  |  |  |  |
| Infants | 0.22 (0.35) | 0.12 (0.15) | 0.18 (0.16) | 0.14 (0.14) | 0.16 (0.17) |
| Children | 0.11 (0.08) | 0.13 (0.08) | 0.19 (0.17) | 0.20 (0.13) | 0.13 (0.11) |
| Adults | 0.09 (0.06) | 0.10 (0.04) | 0.13 (0.13) | 0.15 (0.09) | 0.09 (0.06) |
